# Comprehensive mapping of identical epitopes across human proteins reveals implications for immune recognition and therapeutic design

**DOI:** 10.1101/2025.08.31.673368

**Authors:** Guillaume Kellermann, Olivier Croce, Baharia Mograbi, Paul Hofman, Patrick Brest

## Abstract

Shared epitopes create safety and efficacy issues for T-cell immunotherapy. In order to facilitate the monitoring of immune responses and the engineering required to solve this problem, we performed a computational proteome-wide epitope screening to establish the complete atlas of shared epitopes in the human and murine proteomes. Unlike bacterial or viral antigens, self-antigens like tumor-associated antigens (TAAs) frequently contained a high level of shared MHC-II epitopes identical to unintended other self-proteins. Therefore, shared epitopes should be a mandatory and systematic concern in studies using TAA. Noticeably, a subset of TAAs identified in this atlas is free of this drawback. Therefore, this dataset will be essential for immunologists designing cancer vaccines, but also to interpret immunomonitoring studies against self-antigens in oncology and autoimmunity. To facilitate the detection of common epitopes, a web server has been made available at https://epitopscanner.ircan.org/.

## Introduction

T-cell based immunotherapies, including cancer vaccines and adoptive T-cell receptor (TCR) therapies, rely on the precise recognition of target antigens to achieve therapeutic efficacy while minimizing adverse effects. The success of these approaches depends critically on the specificity of the intended targets, while minimizing off-targets. Past efforts to ensure the safety of immunotherapies have mainly been focused on predicting cross-reactivity for different epitopes^1^, however the obvious problem created by identical sequences shared by different proteins have been neglected^2^. Indeed, two proteins that contain an identical peptide fragment can present an identical epitope to the immune system, leading to off-target autoimmune toxicity. Moreover, the clinical significance of identical shared epitopes extends beyond safety concerns. Preexisting regulatory T-cells recognizing shared epitopes may contribute in explaining the partial tolerance to many tumor-associated antigens (TAA) compared to foreign antigens^3,4^. Finally, shared epitopes complicate the interpretation of immunomonitoring studies, where T-cell responses attributed to specific targets may in fact reflect other unintended proteins.

We have recently emphasized the presence of shared epitopes in several cancer vaccines currently being tested in humans^2^, therefore we realized the need to provide an exhaustive evaluation of this neglected phenomenon for all the antigens of the proteome. To overcome the limitations of current search tools^1^, we have developed a computational approach to list all the shared epitopes from a proteome. Our analysis of the human proteome reveals striking differences between self and foreign antigens (used in vaccines for infectious disease), identifies TAAs with unique epitope profiles suitable for immunotherapy, and provides mechanistic insights into immune tolerance evolution.

## Results

### Comprehensive mapping of all shared identical epitopes

Shared epitopes can be identified using NCBI or UNIPROT BLAST^1^. However, these traditional sequence alignment tools, designed to identify homology between full-length proteins, face significant limitations when trying to list all the short identical peptide sequences within full proteomes. Indeed, these tools fail to detect short amino acid sequences that have a low E-score, thus missing some epitopes. In order to perform an exhaustive listing of all possible shared epitopes, we developed Shared EpitopeMapper, a specialized algorithm designed to systematically scan entire proteomes, including all documented splice variants, specifically searching for redundant sequences of 8 and 11 amino acids: https://epitopscanner.ircan.org/. These length correspond to the minimal epitopes required for presentation by MHC-I and MHC-II molecules respectively^5^, the cellular arm of adaptive immunity.

Our systematic analysis of the human proteome generated two complementary datasets that illuminate the complex landscape of epitope redundancy. First, we catalogued for each gene all the other genes that contain identical MHC epitope sequences, whether 8-mers or 11-mers (Supplementary Tables 1-2). Second, we approached the problem from the opposite angle, documenting for each shared epitope sequence, the complete list of genes harboring that particular sequence (Supplementary Tables 3-4). This dual perspective provides both gene-centric and epitope-centric views of immune reactivity potential.

**Table 1.**

| Enrichment FDR | nGenes | Pathway Genes | Fold Enrichment | Pathways |
| --- | --- | --- | --- | --- |
| 6.7E-38 | 850 | 1830 | 1.4 | Mitochondrion |
| 3.9E-27 | 291 | 520 | 1.7 | Mitochondrial matrix |
| 1.2E-24 | 628 | 1380 | 1.4 | Organelle envelope |
| 1.2E-24 | 628 | 1380 | 1.4 | Envelope |
| 1.5E-23 | 429 | 879 | 1.5 | Mitochondrial envelope |
| 3.7E-23 | 297 | 559 | 1.7 | Mitochondrial inner membrane |
| 2.9E-22 | 324 | 630 | 1.6 | Organelle inner membrane |
| 1.4E-21 | 198 | 339 | 1.8 | Mitochondrial protein-containing complex |
| 9.5E-21 | 399 | 828 | 1.5 | Mitochondrial membrane |
| 2.6E-15 | 74 | 101 | 2.3 | Organelar ribosome |

The analysis identified 557,930 distinct 8-mer sequences and 421,646 distinct 11-mer sequences that are shared between at least two different proteins. The distribution of sharing frequency shows that the top 100 most widely shared 8-mers each occur in more than 68 different genes, while even extending to the top 10,000 8-mers reveals sequences shared by at least 9 different genes (Figure 1A). For longer sequences, the 100 most common 11-mers each appear in over 35 different genes, and the top 10,000 11-mers are each found in at least 8 genes (Figure 1B).

**Figure 1.**
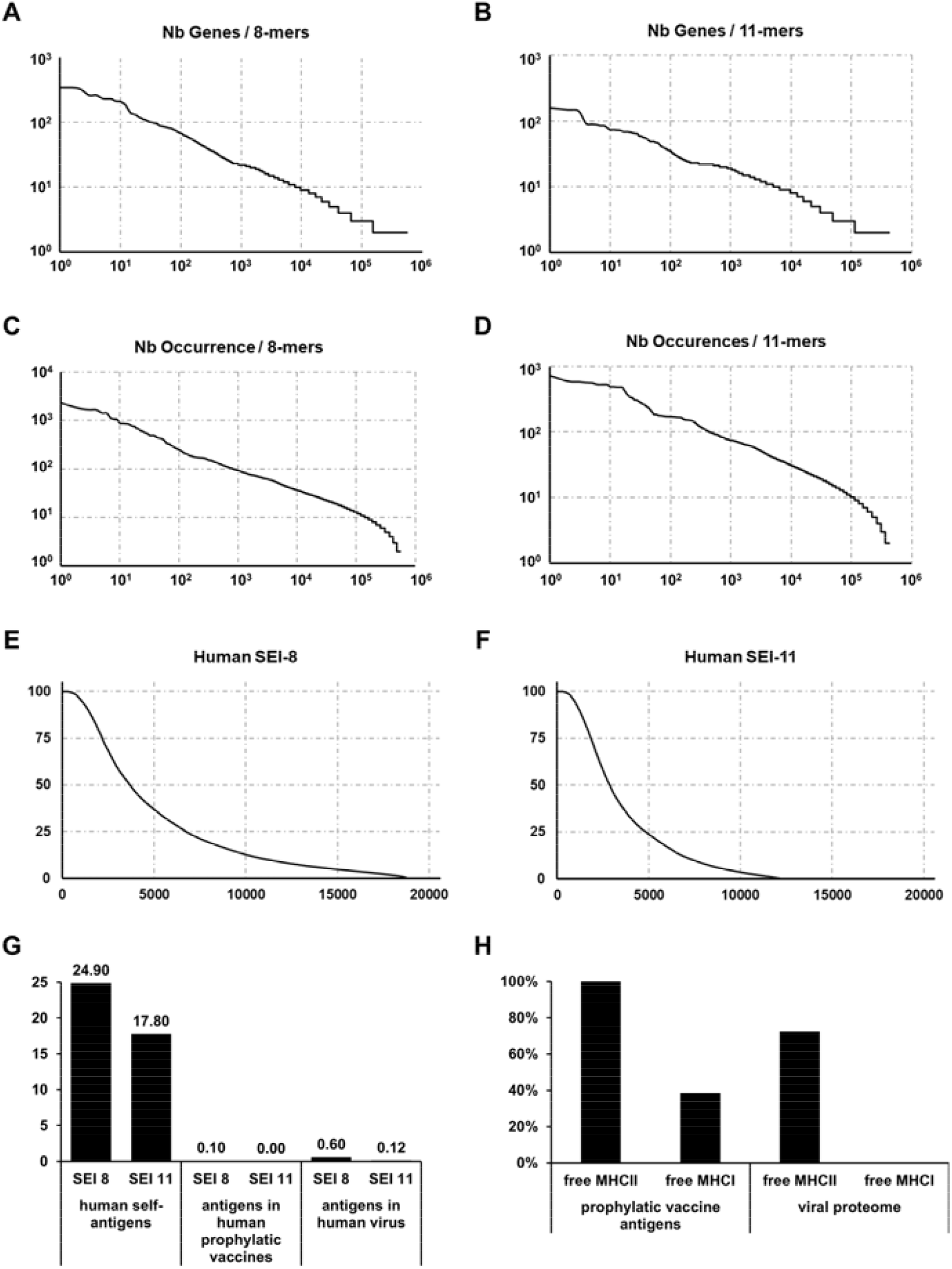
Shared Epitopes Index in proteomes. A) Repartition of genes sharing an identical 8-mers or B) 11-mers in the human proteome. C) Number of occurrence of the shared 8-mers and D) shared 11-mers in the human proteome. E) Distribution of SEI-8 and F) SEI-11 for all human genes across the human proteome. G) comparing the mean SEI of a self-antigen in mammalian proteomes and the mean SEI of foreign (bacterial or viral) antigens with the human proteome (Supplementary 9). H) % of prophylactic vaccines or viral proteomes free of shared MHC-II or MHC-I epitopes.

To account for proteins containing internal repetitive sequences, we calculated the total occurrence frequency of each sequence across the entire proteome. The 100 most abundant 8-mer sequences appear more than 250 times each throughout the human proteome, while the corresponding 11-mers occur more than 171 times each (Figure 1C-D). This analysis captures both inter-protein sharing (sequences found in different proteins) and intra-protein repeats (sequences occurring multiple times within the same protein), thus revealing very complex patterns to consider when trying to predict epitope abundance from gene expression, proteasomal processing, and presentation through the MHC.

### The Shared Epitope Index

To quantify these observations, we introduce the Shared Epitope Index (SEI), defined as the percentage of epitopes within a protein that are also found in other proteins. We calculated SEI-8 and SEI-11 separately to account for the different requirements of MHC-I and MHC-II presentation pathways (Figure 1E-F).

The proteome-wide averages were 24.9% for SEI-8 and 17.8% for SEI-11, indicating that nearly one-quarter of all potential 8-mer epitopes in human proteins are shared with other proteins. Our analysis also found that 13.9% of all human genes are composed predominantly of shared 11-mer sequences. Moreover, some proteins reached 100% SEI-11, meaning that every possible 11-amino acid segment within these proteins can also be found elsewhere in the proteome.

Conversely, 8.7% of human genes contained no shared 8-mer epitopes, and 41% contained no shared 11-mers (Figure 1E-F). The gene ontology of the proteins with limited epitope sharing revealed an enrichment for mitochondrial genes, particularly those encoding components of the respiratory chain complexes (Table 1). This observation indicates a lower frequency of orphan genes in nuclear versus mitochondrial genomes.

### Antigens used in prophylactic vaccines against infectious disease share little overlap with human proteome

To contextualize the high SEI values of self-antigens, we separately searched for shared epitopes with the human proteome, but now in the antigens used in prophylactic vaccines to prevent infectious disease. Across 13 foreign antigens from licensed prophylactic vaccines, we identified 18 8-mers that were identical to sequences in the human proteome, representing a total mean of 0.1% SEI-8 and 0% SEI-11. Then, we extended this comparison to the proteome of 29 human virus and found only a mean of 0.6% for SEI-8 and 0.12% for SEI-11 (Supplementary Table 5). These analyses emphasize a striking difference between foreign and self-antigens (Figure 1G). *Foreign antigens exhibit minimal sequence overlap with the human proteome, while self-antigens often share a large proportion of epitopes*.

All virus and some licensed vaccine antigens contained a low content (<2%) of MHC-I epitopes, whereas all vaccines antigens and most virus are free of shared MHC-II epitopes (Figure 1H). The shared MHC-II epitopes detected are often limited to low-complexity repetitive sequences in large DNA viruses (Supplementary 6). Therefore, while a low content of shared MHC-I epitopes is commonly encountered in foreign antigens, shared MHC-II epitopes are much rarer. This provides a rational to explain why CD4 T-cells, but not CD8 T-cells, have been evolutionary selected to play the leading role in the regulation of immune tolerance. Shared MHC-II epitopes are usually absent in foreign antigens, preventing their recognition by preexisting Tregs, and thus supporting their immunogenicity, contrary to self-proteins.

### Shared MHC-II epitopes are frequent in TAA

In stark contrast to human viruses and prophylactic vaccine antigens, shared MHC-II epitopes are abundant in the majority of the common TAAs used in cancer vaccines (Figure 2A). In some cases, shared MHC-II epitopes accounted for the major portion of the antigen. This extensive sequence overlap reduces the target specificity of many TAAs and may create off-target issues. The POTEE tumor antigen exemplifies this issue, as it is composed of 99.08% of shared 11-mers. Similarly, more than 95% of the sequence of MAGE-A3 is covered by partially overlapping fragments from other MAGE proteins. Consequently, the only unique MHC epitopes for MAGE-A3 are restricted to the sequences covering the junctions between the shared sequences and are representing only 4.7% of all the possible 11-mer peptides from this gene (Figure 2B). These high SEI level show that the problem of shared epitope should be a mandatory and systematic concern when using TAA, because recent evolution by gene duplication has led to extensive sequence identity across the proteome.

**Figure 2.**
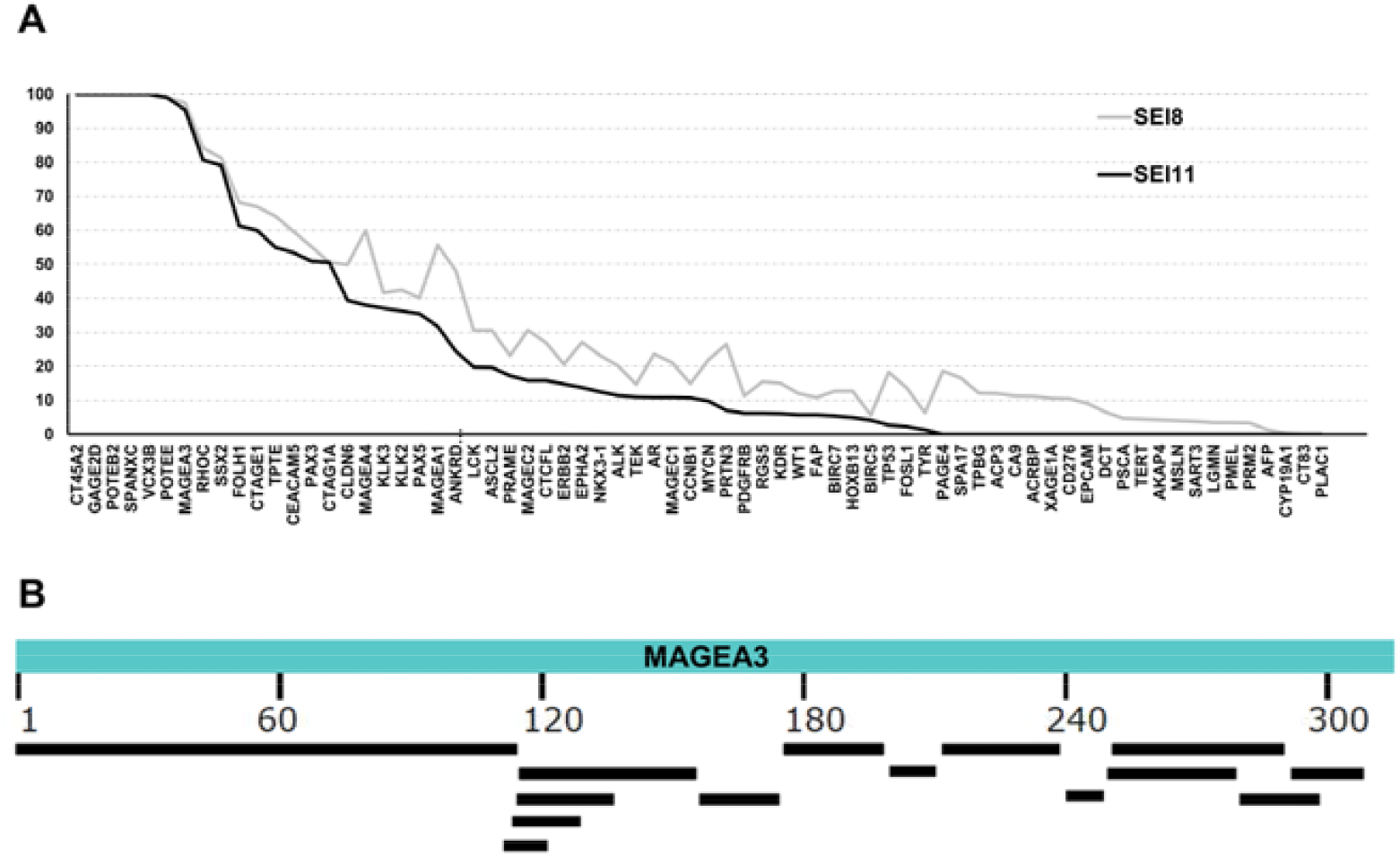
Shared Epitopes in TAA. A) SEI-8 and SEI-11 in common TAA used in therapeutic cancer vaccines. B) Alignments of shared sequences in MAGEA3.

Interestingly, we identified a subset of TAAs that is free of shared MHC-II epitope and, in some cases, do not contain any shared MHC-I epitope with the human proteome (Figure 2A). KK-LC-1, PLAC1, PSCA, PMEL-gp100, CYP19A1, AFP, SART3, CD276, MSLN, AKAP4, TERT, XAGE1, PRM2, LGMN and ACRBP emerge from this atlas as these antigens exhibit a high degree of sequence uniqueness and provide unambiguous targets for cancer immunotherapies.

### Relevance of mouse models to study shared epitopes

By comparing our exhaustive catalog of shared epitopes with mouse orthologs, we found that the SEI of a human gene is usually very close to its corresponding ortholog in mice (supplementary Tables 6-9), especially when the SEI is low (Figure 3A-B). However, this comparable SEI is mostly due to parallel evolution, as only 41% of shared 8-mer epitopes are identical between both species (Figure 3C), indicating species-specific differences in the actual epitope sequences despite similar shared patterns. These observations show that mouse models will be limited for evaluating the safety of shared epitopes from human antigens. *In conclusion, the effects of shared epitopes may often be modelized in mice, but only with the use of mouse antigens*.

**Figure 3.**
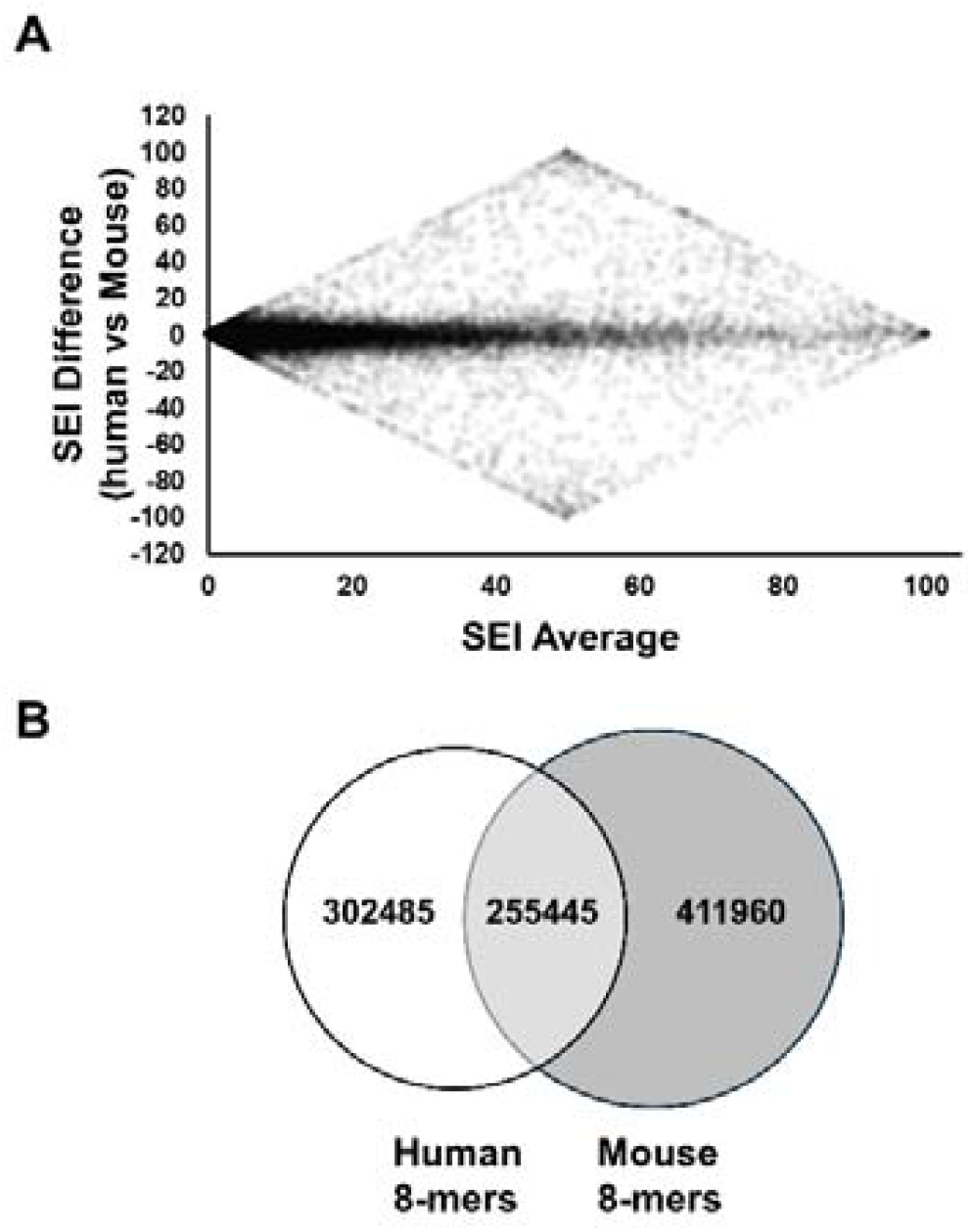
Differential SEI-8 in mice and human. A) Bland Altman plot to analyze the agreement in SEI-8 between the 17013 genes in common between these two species. B) Venn diagram to compare the shared and unique epitopes between these two species.

## Discussion

Our comprehensive proteome-wide epitope analysis reveals a fundamental difference between the foreign and the self-antigens used in vaccines. Prophylactic foreign antigens are free of MHC-II epitopes shared with human proteins, whereas TAA contain extensive sequences that overlap with endogenous, non-targeted proteins. This high level of epitope sharing necessary impairs both TAA immunogenicity and their tumor specificity.

Even oncofetal TAAs with restricted tumor expression, such as POTEE^6^, lose their specificity because they share epitopes with other proteins. Pre-existing CD4+ regulatory T-cells that may recognize these shared epitopes provide a mechanistic explanation for the partial tolerance often observed to oncofetal TAA^3,4^. Recognition of the importance of SEI now calls for a reassessment of the past TAA prioritization rank for cancer immunotherapy^7^. Among the commonly studied TAAs, KK-LC-1, TERT, PLAC1, CYP19A1, AFP, AKAP4, XAGE1, PRM2 and ACRBP emerge as superior candidates due to their high content of unique epitopes and limited somatic expression. These TAAs exhibit greater “foreignness” compared to conventional TAAs and probably also many neoantigens^8^. Alternatively, our comprehensive shared epitope catalog will facilitate the systematic removal of problematic sequences to create reengineered TAAs for the next-generation of cancer vaccines.

Identification of shared epitopes is also critical for an accurate immunomonitoring. For example, the T-cell responses against MAGE-A3_212-220_ after BNT111 vaccination^9^ provides an example of this challenge: this epitope is shared with MAGE-A12, raising safety concerns regarding off-target neurotoxicity^10^, but also suggesting potential therapeutic advantages through a broader TAA coverage. Strategic targeting of epitopes shared by multiple TAAs could indeed improve therapeutic coverage, reduce treatment failures due to the loss of single antigens, while expanding patient selection beyond traditional single-determinant criteria. Multiple antigens with modest individual overexpression could cumulatively share epitopes at therapeutically relevant, differential levels in tumors, if multiple antigens provide the same shared epitopes, ultimately creating valid targets in patients that are currently missed by inclusion criteria that do not include shared epitopes in their analysis.

In conclusion, the catalogs of genes and shared epitopes together with the web server provided in this work represent the definitive resource for the systematic identification of all targets of MHC epitopes used in immunological studies. Therefore, this study opens the way to a better understanding of the risks and benefits of shared epitopes in cancer immunotherapy and autoimmunity.

## Material and Methods

We developed an **open-source suite of Python scripts** (EpitopeMapper, KmerProteomeComparator□ and EpitopeCrossScanner) to enumerate k-mers, compare proteomes and identify shared epitopes. The full code and its full documentation are freely available at https://github.com/IRCAN/EpitopeMapper (login: epitopemapper-test password: #epitopemapper-test2025)

### Proteome data sets

Reviewed human (UP000005640) and mouse (UP000000589) reference proteomes, including all canonical and isoform sequences, were downloaded from UniProt release□2025□_02 (16□April□2025). **Human proteome includes:** □58□520 protein sequences (20□396 canonical□+□38□124 isoforms; 32□418□746 amino-acid residues) and mouse proteome includes 41□489 protein sequences (22□568 canonical□+□18□921 isoforms; 23□764□281 residues). In addition, sequences of 13 licensed vaccine antigens and of 29 human viral proteomes were retrieved from UniProt on the same date (Supplementary□Table□5). For each antigen the gene-product FASTA file was used as supplied by UniProt.

### K-mers enumeration and indexation

K-mers were searched using a sliding window of size **k**□**=**□**8** (minimal MHC-I) or **k**□**=**□**11** (MHC-II 9-mer core□+□2 peptide-flanking residues)^5^, every contiguous peptide was extracted with step□=□1. K-mers containing a masked residue (ie.: **B, J, O, U, X, Z)** were excluded.

For each k-mer we stored the parent gene symbol, UniProt accession, positional coordinates and an isoform flag.

### Definition of the Shared Epitope Index (SEI)

For a gene *g* and k-mer length *k*, let

*P*_*g,k*_□=□set of distinct k-mers encoded by *g*

*P*_*¬g,k*_□=□union of k-mers encoded by all other genes.

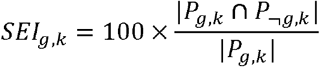

SEI-8 and SEI-11 were computed independently for every human and mouse gene. Genes with SEI□=□0 were classified as epitope-unique.

### Cross-species k-mer comparison

To quantify conservation of potential epitopes between species we used the home made script **KmerProteomeComparator**, which (i) extracts all k-mers from two proteomes, (ii) partitions them into *shared, only-in-proteome*□*1*, and *only-in-proteome*□*2* sets, and (iii) records absolute and relative counts Pairwise human-versus-mouse analyses were performed for k□=□8 and k□=□11.

### Detection of shared epitopes between foreign antigens and the human proteome (EpitopeCrossScanner v0.3)

Each vaccine antigen or viral protein was scanned with the same sliding-window algorithm (script EpitopeCrossScanner.py). Identical 8-mer and 11-mer matches in the human proteome were recorded, and SEI values for foreign antigens were calculated analogously to the procedure above.

## Author contribution

GK and PB performed Conceptualization, Writing - Original Draft, Visualization. Funding acquisition; OC performed Methodology. PH and BM performed Writing - Review & Editing and Funding acquisition.

## Conflict of interest

GK is shareholder of Telomium, a company developing cancer vaccines.

## Funding

The author(s) declare financial support was received for the research, authorship, and/or publication of this article. Funding is acknowledged from the French Government (Agence Nationale de Recherche, ANR) through RNPVAX (ANR-23-CE17-0011-01, ANR-23-CE17-0011-03), “IHU RespirERA” France 2030 program (ANR-23-IAHU-007), “Investments for the Future” programs LABEX SIGNALIFE ANR-11-LABX-0028-01, IDEX UCAJedi ANR-15-IDEX-01, Canceropole PACA; Région Sud, CNRS, INSERM cancer; INCA Plan Cancer; ITMO Cancer.

## Legend of figure

**Supplementary Table 1**. List of genes from the human proteome sorted for their content in shared MHC epitopes (11aa).

**Supplementary Table 2**. List of genes from the human proteome sorted for their content in shared MHC epitopes (8aa).

**Supplementary Table 3**. List of shared 11-mers from the human proteome.

**Supplementary Table 4**. List of shared 8-mers from the human proteome.

**Supplementary Table 5**. shared 8-11mers with the human proteome in prophylactic vaccines or viral proteomes.

**Supplementary Table 6**. List of genes from the murine proteome sorted for their content in shared MHC epitopes (11aa).

**Supplementary Table 7**. List of genes from the murine proteome sorted for their content in shared MHC epitopes (8aa).

**Supplementary Table 8**. List of shared 11-mers from the murine proteome.

**Supplementary Table 9**. List of shared 8-mers from the murine proteome.

## Footnotes

1 https://blast.ncbi.nlm.nih.gov/Blast.cgi https://www.uniprot.org/blast

## Notes

https://epitopscanner.ircan.org/

## References

1. Jaravine, V., Raffegerst, S., Schendel, D. J. & Frishman, D. Assessment of cancer and virus antigens for cross-reactivity in human tissues. Bioinforma. Oxf. Engl. 33, 104–111 (2017).

2. Kellermann, G., Mograbi, B., Hofman, P. & Brest, P. Shared epitopes create safety and efficacy concerns in several cancer vaccines. J. Immunother. Cancer 13, (2025).

3. Greilach, S. A. et al. Presentation of Human Neural Stem Cell Antigens Drives Regulatory T cell induction. J. Immunol. Baltim. Md 1950 210, 1677–1686 (2023).

4. Huijbers, I. J. et al. Minimal tolerance to a tumor antigen encoded by a cancer-germline gene. J. Immunol. Baltim. Md 1950 188, 111–121 (2012).

5. Arnold, P. Y. et al. The majority of immunogenic epitopes generate CD4+ T cells that are dependent on MHC class II-bound peptide-flanking residues. J. Immunol. Baltim. Md 1950 169, 739–749 (2002).

6. Bassoy, E. Y. et al. Identification of TTLL8, POTEE, and PKMYT1 as immunogenic cancer-associated antigens and potential immunotherapy targets in ovarian cancer. OncoImmunology 14, 2460276 (2025).

7. Cheever, M. A. et al. The Prioritization of Cancer Antigens: A National Cancer Institute Pilot Project for the Acceleration of Translational Research. Clin. Cancer Res. Off. J. Am. Assoc. Cancer Res. 15, 5323–5337 (2009).

8. Singhaviranon, S., Dempsey, J. P., Ebrahimi-Nik, H. & Srivastava, P. K. Defining key parameters of therapeutic activity using mouse cancer neoepitopes. J. Immunol. vkaf090 (2025) doi:10.1093/jimmun/vkaf090.

9. Sahin, U. et al. An RNA vaccine drives immunity in checkpoint-inhibitor-treated melanoma. Nature 585, (2020).

10. Morgan, R. A. Cancer regression and neurological toxicity following anti-MAGE-A3 TCR gene therapy. J Immunother Hagerstown Md 36, 133–151 (2013).

